# Pre-therapeutic Microglia Activation and Sex Determine Therapy Effects of Chronic Immunomodulation

**DOI:** 10.1101/2021.05.30.445761

**Authors:** Gloria Biechele, Tanja Blume, Maximilian Deussing, Benedikt Zott, Yuan Shi, Xianyuan Xiang, Nicolai Franzmeier, Gernot Kleinberger, Finn Peters, Katharina Ochs, Carola Focke, Christian Sacher, Karin Wind, Claudio Schmidt, Simon Lindner, Franz-Josef Gildehaus, Florian Eckenweber, Leonie Beyer, Barbara von Ungern-Sternberg, Peter Bartenstein, Karlheinz Baumann, Mario M. Dorostkar, Axel Rominger, Paul Cumming, Michael Willem, Helmuth Adelsberger, Jochen Herms, Matthias Brendel

**Author notes:** **Corresponding author:** Dr. Matthias Brendel, MHBA, Department of Nuclear Medicine, University Hospital of Munich, Marchioninistr.15, 81377 Munich, Germany.

## Abstract

Modulation of the innate immune system is emerging as a promising therapeutic strategy against Alzheimer’s disease (AD). However, determinants of a beneficial therapeutic effect are ill-understood. Thus, we investigated the potential of 18 kDa translocator protein positron-emission-tomography (TSPO-PET) for assessment of microglial activation in mouse brain before and during chronic immunomodulation. Serial TSPO-PET was performed during five months of chronic microglia modulation by stimulation of peroxisome proliferator-activated receptor (PPAR)-γ with pioglitazone in two different mouse models of AD (PS2APP, *App^NL-G-F^*). Using mixed statistical models on longitudinal TSPO-PET data, we tested for effects of therapy and sex on treatment response. We tested correlations of baseline with longitudinal measures of TSPO-PET, and correlations between PET results with spatial learning performance and β-amyloid accumulation of individual mice. Immunohistochemistry was used to determine the molecular source of the TSPO-PET signal. Pioglitazone-treated female PS2APP and *App^NL-G-F^* mice showed attenuation of the longitudinal increases in TSPO-PET signal when compared to vehicle controls, whereas treated male *App^NL-G-F^* mice showed the opposite effect. Baseline TSPO-PET strongly predicted changes in microglial activation in treated mice (R=−0.874, p<0.0001) but not in vehicle controls (R=−0.356, p=0.081). Reduced TSPO-PET signal upon treatment was associated with better spatial learning and higher fibrillar β-amyloid accumulation. Immunohistochemistry confirmed activated microglia to be the source of the TSPO-PET signal (R=0.952, p<0.0001). TSPO-PET represents a sensitive biomarker for monitoring of immunomodulation and closely reflects activated microglia. Pre-therapeutic assessment of baseline microglial activation and sex are strong predictors of individual immunomodulation effects and could serve for responder stratification.

## Introduction

Neuroinflammation is now recognized as an inherent part of the Alzheimer’s disease (AD) pathology ^1^. The key players of neuroinflammation in AD are activated microglia and astrocytes ^2^. Although it is still unclear if beneficial or detrimental effects of neuroinflammation dominate in the (patho)physiology of AD, there is considerable interest in integrating the modulation of neuroinflammation into novel treatment strategies against AD ^3^. Preclinical studies showed that immunomodulation by peroxisome proliferator-activated receptor (PPAR)-γ using the antidiabetic compound pioglitazone rescues neuronal spine density ^4^ and spatial learning performance ^5^ in mouse AD models. However, a large human trial with pioglitazone in mild cognitive impairment due to AD was terminated after an interim analysis showing lack of efficacy ^6^. Hence, the discrepancies between beneficial effects in preclinical studies and lacking efficacy in humans deserve detailed inquiry, with the objective of uncovering the salient factors accounting for the failure of PPARγ stimulation in clinical translation.

TSPO-PET is increasingly used to monitor therapy-related changes of microglial activation in humans ^7^ and rodent models ^8^. In this regard, the TSPO ligand ^18^F-GE180 is proven effective for robust imaging of microglial activation in a mouse model of amyloidosis, and the normalization of TSPO binding upon treatment with neurotrophin receptor ligand that ameliorates hyperphosphorylation and misfolding of tau and rescues the consequent neurite degeneration ^9^. Our previous data revealed excellent agreement between ^18^F-GE180 PET quantitation and immunohistochemistry of microglial markers ^10, 11^, thus indicating its potential to access and predict PPARγ stimulation effects *in vivo*.

Therefore, we aimed to test the hypothesis that TSPO-PET with ^18^F-GE180 is a suitable tool for monitoring anti-neuroinflammatory responses to chronic immunomodulation in AD mouse models. We furthermore tested the hypothesis that microglial activation by TSPO-PET predicts therapy related changes and outcome parameters. Furthermore, we tested for effects of mouse sex on immunomodulation. Finally, we used immunohistochemistry to validate *in vivo* PET findings and to confirm the cellular source of TSPO-PET signal alterations.

## Results

### TSPO-PET detects altered microglia activation during chronic PPARγ stimulation

First, we investigated whether effects of chronic PPARγ stimulation can be detected by TSPO PET in PS2APP mice and wild-type controls. Vehicle-treated PS2APP mice showed a strong increase over time of the TSPO-PET signal when compared to vehicle-treated wild-type mice between eight and 13 months of age, with a peak at 11.5 months (+52-67%, all time-points: p < 0.0001, **Figure 1**). The pre-therapeutic baseline TSPO-PET signal did not significantly differ between PS2APP mice with and without pioglitazone treatment (SUV_H_: 0.24±0.05 vs. 0.26±0.01, p = 0.647). However, PS2APP mice with pioglitazone treatment had a lower TSPO-PET signal at 9.5 (−13%, p = 0.0027), 11.5 (−17%, p = 0.0046), and 13.0 (−13%, p = 0.0071) months of age when compared to age-matched vehicle-treated PS2APP mice (**Figure 1**). Linear mixed models revealed a main effect of treatment group on TSPO-PET across time-points (b/SE=−0.036/0.006, T=5.405, p<0.0001), controlling for age (i.e. quadratic effect) and random intercept. Individual PS2APP mice indicated a heterogeneous pharmacotherapy-related change in the TSPO-PET signal, which was already conspicuous during the first six weeks of treatment (range of change: −35% to +86%). Pioglitazone-treated wild-type mice manifested a slight decrease of the TSPO-PET signal after six weeks of treatment when compared to vehicle-treated wild-type mice (−12%, p = 0.013), and no such differences at the other time points. Taken together, we found strong differences of TSPO expression *in vivo* between treated and non-treated animals, suggesting the feasibility of TSPO-PET to monitor effects of PPARγ stimulation.

**Figure 1.**
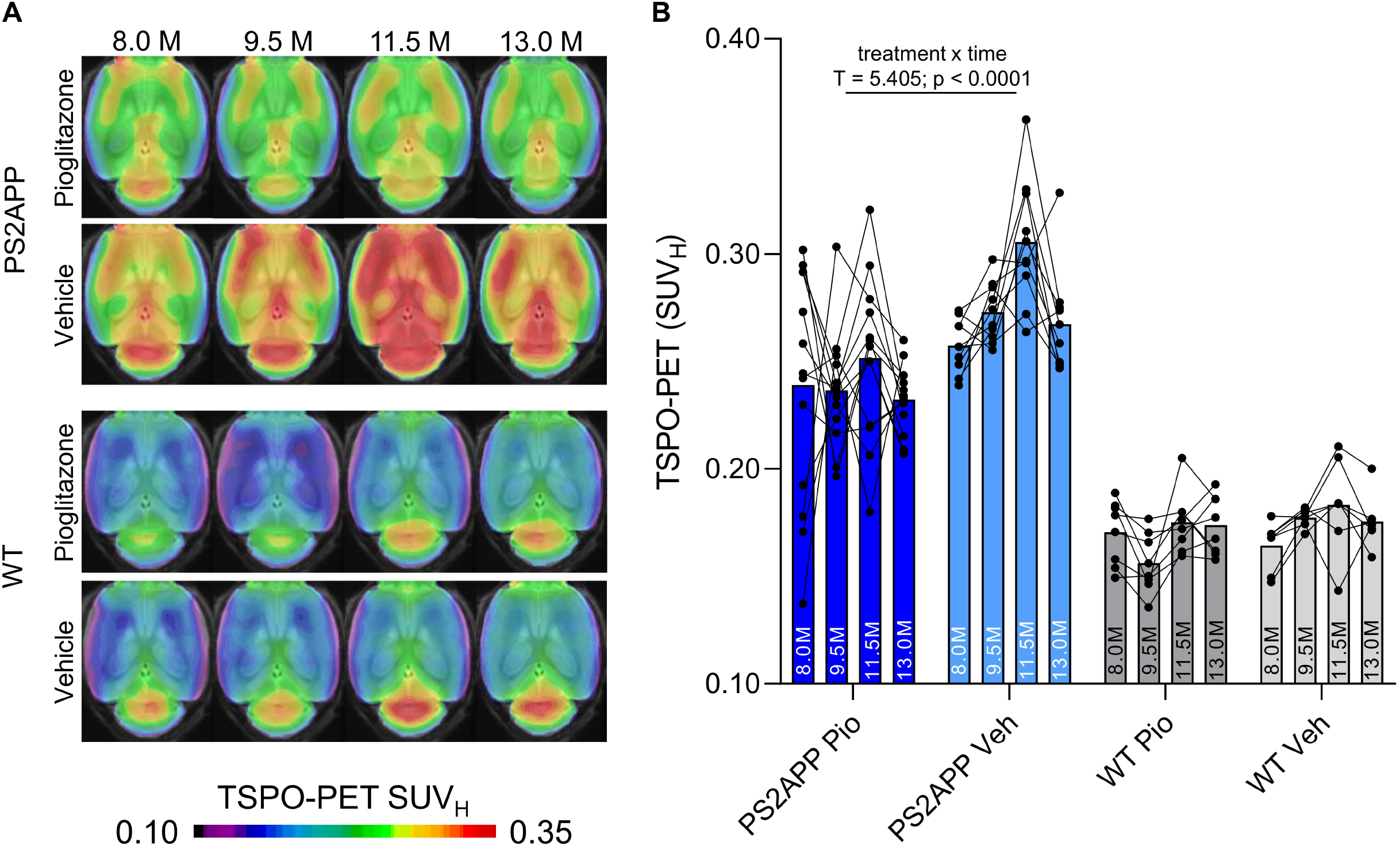
TSPO-PET monitoring of chronic pioglitazone treatment in PS2APP and wild-type (WT) mice. (**A**) Axial images show group levels of the ^18^F-GE180 TSPO-PET signal (myocardium scaled standardized uptake value, SUV_H_) at different ages in treatment and vehicle groups, projected upon a standard MRI anatomic template. Baseline scans were performed prior to treatment initiation. (**B**) Individual time courses of the cortical TSPO-PET signal during the treatment period. Pio = pioglitazone treatment, Veh = vehicle treatment. Statistics derive from a linear mixed model. PS2APP pioglitazone n=13, PS2APP vehicle n=10, WT pioglitazone n=8, WT vehicle n=7.

### Chronic PPARγ stimulation changes microglial activation independent of APP overexpression but dependent on sex

Next, we tested whether previously observed sex differences in TSPO expression in mouse brain ^12^ have an impact on the responses to PPARγ pharmacological stimulation. To this end, we used the novel APP knock-in model *App^NL-G-F^* ^13^ mice and performed longitudinal TSPO-PET imaging during chronic pioglitazone treatment in groups of female and male mice. Furthermore, we tested whether these mice showed effects of PPARγ stimulation on the TSPO-PET signal in the absence of APP overexpression. We observed sex-specific elevation of the TSPO-PET signal in vehicle-treated female *App^NL-G-F^* mice when compared to males aged 7.5 (+18%, p = 0.017) and 10 months of age (+25%, p = 0.0007; sex x time interaction: b/SE=−0.100/0.030, T = −3.273, p = 0.0003, linear mixed model controlling for random intercept). Female *App^NL-G-F^* mice with pioglitazone treatment showed normalization of the TSPO-PET signal increase compared to vehicle-treated female *App^NL-G-F^* mice aged from five to ten months (**Figure 2**). This resulted from decreased TSPO-PET signal in pioglitazone-treated female *App^NL-G-F^* mice aged 7.5 (−15%, p = 0.030) and 10 months (−21%, p = 0.0053) when compared to vehicle-treated female *App^NL-G-F^* mice (treatment x time interaction: b/SE=0.114/0.030, T=3.801, p=0.0009, linear mixed model controlling for random intercept, **Figure 2**). On the other hand, male *App^NL-G-F^* mice with pioglitazone treatment tended to show an exaggerated increase of the TSPO-PET signal from five to ten months of age when compared to vehicle treated male *App^NL-G-F^* mice (+12% vs. +2%, treatment x time interaction: b/SE=−0.041/0.022, T=−1.862, p=0.072). Wild-type mice did not show differences in TSPO-PET signal between treated or non-treated animals for this time span. In summary, sex strongly influenced the magnitude of microglial response to amyloidosis, with distinctly opposite effects in female and male mice. APP overexpression at baseline was not a significant determinant of effects of PPARγ stimulation on TSPO expression.

**Figure 2.**
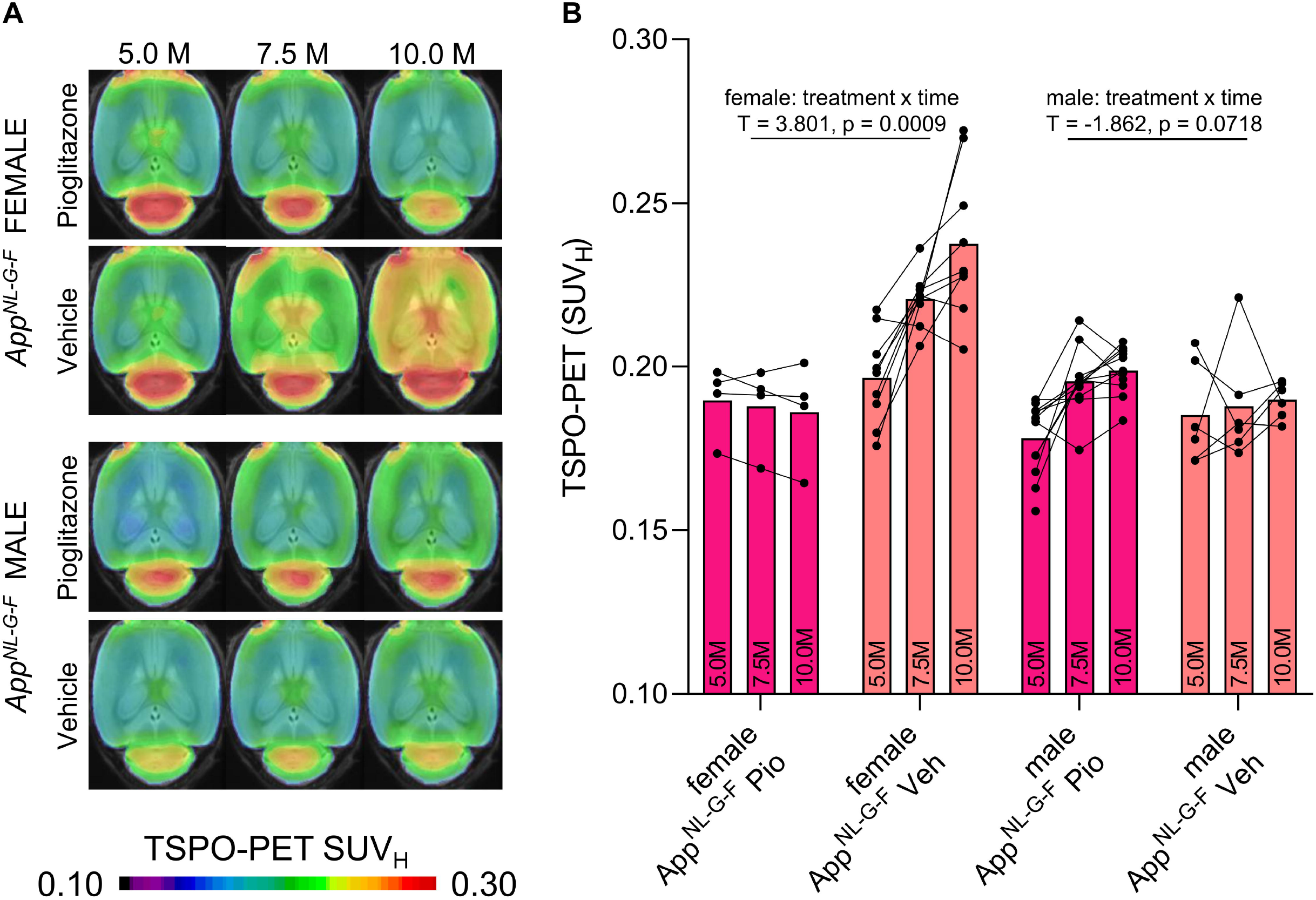
^18^F-GE180 TSPO-PET monitoring of effects of chronic pioglitazone treatment in female and male *App^NL-G-F^* mice. (**A**) Axial images show group means of the TSPO-PET signal (myocardium scaled standardized uptake value, SUV_H_) at different ages in treatment and vehicle groups, projected upon a standard MRI template. Baseline scans were performed prior to therapy initiation. (**B**) Individual time courses of the cortical TSPO-PET signal during the pharmacological treatment period. Statistics derive from a linear mixed model. Female *App^NL-G-F^* pioglitazone n=4, female *App^NL-G-F^* vehicle n=9, male *App^NL-G-F^* pioglitazone n=11, male *App^NL-G-F^* vehicle n=6. Pio = pioglitazone treatment, Veh = vehicle treatment.

### Baseline TSPO-PET predicts treatment associated changes in microglial activation during chronic PPARγ stimulation

Given the observed heterogeneity of changes in TSPO-PET after induction of PPARγ stimulation, we asked if TSPO-PET at baseline serves to predict the individual longitudinal changes in microglial activation upon treatment. Strikingly, we observed a strong negative association between baseline TSPO-PET and subsequent changes in the TSPO-PET signal across pioglitazone treated animals (R = −0.874, p < 0.001, **Figure 3A, B**), suggesting that mice with high microglial activation at baseline respond stronger to PPARγ stimulation. Importantly, this association was also present in independent cohorts of PS2APP mice (R = −0.964, p < 0.0001) and *App^NL-G-F^* mice (R = −0.680, p = 0.0053) mice with chronic pioglitazone treatment. On the other hand, there was only a trend towards a negative association between the baseline TSPO-PET signal and subsequent TSPO-PET changes in vehicle treated animals (R = −0.356, p = 0.081). The association between pre-therapeutic TSPO-PET results and changes in microglial activation of pioglitazone treated mice was observed across all brain regions, with the strongest differences relative to vehicle treated mice in neocortical areas, hippocampus, striatum, and thalamus (**Figure 3C, D; Table 1**). Several subcortical regions also showed a significant negative association between the baseline TSPO-PET signal and changes in microglial activation in the vehicle cohort (**Table 1**), congruent with the observation that microglial activation at baseline per se has a predictive value for longitudinal alterations of microglial activity and spatial learning performance ^14, 15^. In summary, our data indicate a strong predictive value of a baseline biomarker of microglial activation for the effect of immunomodulatory therapy.

**Figure 3.**
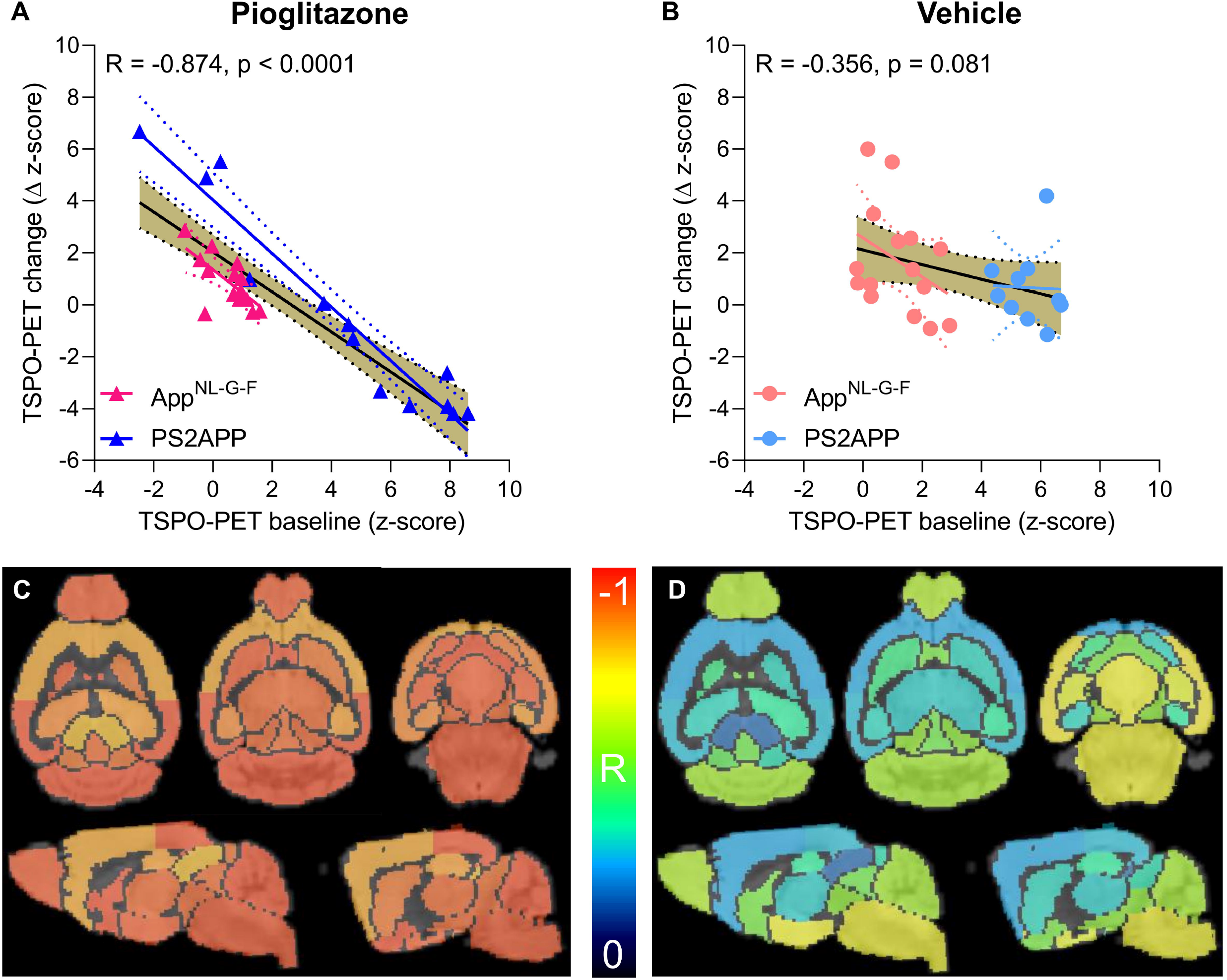
Prediction of changes in microglial activity by the ^18^F-GE180 TSPO-PET baseline examination. (**A, B**) Correlation analysis between the TSPO-PET z-score at baseline and the change of the TSPO-PET z-score (baseline to last follow-up) in *App^NL-G-F^* and PS2APP mice with pioglitazone treatment (**A**) and vehicle controls (**B**). Correlations are illustrated for the combination of both mouse models (gold) and in separate analyses of *App^NL-G-F^* (red) and PS2APP (blue) mice. (**C, D**) Multiregional analysis of the correlation between the TSPO-PET z-score at baseline and the change of the TSPO-PET z-score (baseline to last follow-up). Reprojected coefficients of correlation (R) are illustrated in axial and sagittal slices projected upon a standard MRI template. *App^NL-G-F^* and PS2APP mice were analyzed together for the pioglitazone treatment group (**C**) and vehicle controls (**D**). Levels of significance per region after false discovery rate correction for multiple comparisons are provided in **Table 1**.

**Table 1.** Multi-region analysis of baseline prediction of longitudinal microglial activation by baseline TSPO-PET. Person’s correlation coefficients (R) were calculated between baseline TSPO-PET (z-score) and the change in TSPO-PET (Δ z-score) during the five months treatment period in pioglitazone and vehicle treated *App^NL-G-F^* and PS2APP mice. P-values were adjusted for multiple comparisons by false discovery rate correction. *p<0.05; **p<0.01; ***p<0.001

| Region | Pioglitazone |  | Vehicle |  | Contrast |
| --- | --- | --- | --- | --- | --- |
|  | R | p-value (FDR-corrected) | R | p-value (FDR-corrected) |  |
| Striatum R | -0.941 | 4.9e-13*** | -0.427 | 0.12 | 0.514 |
| Striatum L | -0.899 | 1.0e-10*** | -0.342 | 0.12 | 0.557 |
| Hippocampus R | -0.879 | 8.6e-10*** | -0.344 | 0.12 | 0.535 |
| Hippocampus L | -0.854 | 8.6e-09*** | -0.388 | 0.083 | 0.466 |
| Thalamus | -0.920 | 7.8e-12*** | -0.326 | 0.13 | 0.594 |
| Cerebellum | -0.952 | 1.4e-13*** | -0.584 | 0.0066** | 0.368 |
| Basal forebrain & septum | -0.949 | 1.4e-13*** | -0.516 | 0.017* | 0.433 |
| Hypothalamus | -0.936 | 8.6e-13*** | -0.705 | 0.0009*** | 0.231 |
| Amygdala R | -0.935 | 8.6e-13*** | -0.665 | 0.0012** | 0.270 |
| Amygdala L | -0.935 | 9.0e-13*** | -0.666 | 0.0015** | 0.269 |
| Brainstem | -0.943 | 1.4e-12*** | -0.691 | 0.0009*** | 0.252 |
| Central grey | -0.914 | 1.8e-11*** | -0.505 | 0.019* | 0.409 |
| Superior colliculi | -0.804 | 2.6e-07*** | -0.199 | 0.34 | 0.605 |
| Olfactory bulb | -0.938 | 7.5e-13*** | -0.603 | 0.0050** | 0.335 |
| Midbrain R | -0.919 | 9.5e-12*** | -0.558 | 0.0099** | 0.361 |
| Midbrain L | -0.922 | 6.4e-12*** | -0.541 | 0.012* | 0.381 |
| Inferior colliculus R | -0.900 | 3.0e-11*** | -0.486 | 0.11 | 0.414 |
| Inferior colliculus L | -0.910 | 9.7e-11*** | -0.357 | 0.024* | 0.553 |
| Piriform/entorhinal cortex | -0.883 | 6.2e-10*** | -0.742 | 0.0005*** | 0.141 |
| Auditory/visual cortex | -0.947 | 1.8e-13*** | -0.306 | 0.15 | 0.641 |
| Motor/sensory cortex | -0.836 | 3.2e-08*** | -0.283 | 0.18 | 0.553 |

### PPARγ stimulation induced changes of microglial activation predict spatial learning performance and aggregation of fibrillar Aβ

Next, we asked if altered TSPO expression during chronic pioglitazone treatment has associations with known determinants of therapeutic effects. To this end, we correlated the rate of change in the TSPO-PET signal during the treatment period with the individual spatial learning impairment and changes in fibrillary Aβ pathology measured post mortem. Better spatial learning was associated with an attenuated increase of the TSPO-PET signal during five months of PPARγ stimulation in PS2APP mice (R = −0.733, p = 0.0043, **Figure 4A, B**), but the association did not reach statistical significance in *App^NL-G-F^* mice (R = −0.349, p = 0.221, **Figure 4C, D**). The observed effect in PS2APP mice was treatment-specific, since there was no association between altered TSPO expression and spatial learning in vehicle treated mice (R = −0.032, p = 0.991, **Figure 4B**). Our dedicated analysis of Aβ species during chronic PPARγ stimulation in this same cohort ^16^ revealed a greater increase in fibrillar Aβ, which is the major source of the Aβ-PET signal ^17^, in both treated mouse models compared to their vehicle controls, which reflected a shift of Aβ plaques towards a more fibrillary composition. Meanwhile, the non-fibrillar proportion of plaques decreased upon the treatment, as is reported elsewhere ^16^. A low area under the curve (AUC) of TSPO-PET signal during the recording period was associated with a higher rate of change of fibrillar Aβ in pioglitazone-treated PS2APP (R = −0.600, p = 0.030, **Figure 4E, F**) and *App^NL-G-F^* mice (R = −0.553, p = 0.040, **Figure 4G, H**). Vehicle controls of both models did not show significant associations between the TSPO-PET AUC and changes in fibrillary Aβ pathology. Thus, imaging results for our treatment paradigm confirmed that higher microglial activation is associated with a slower rate of Aβ accumulation, suggesting a net protective effect ^18^.

**Figure 4.**
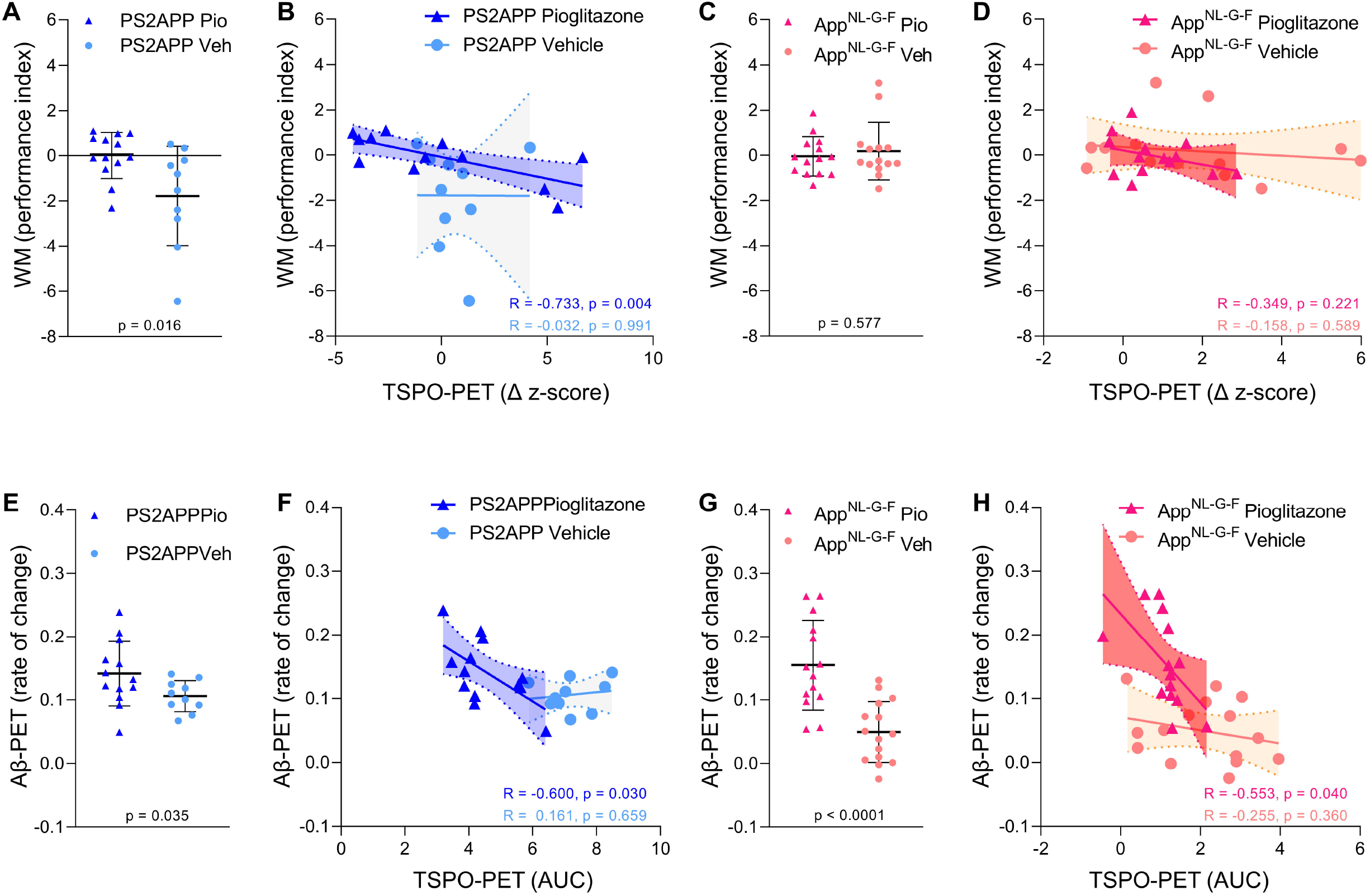
Associations of ^18^F-GE180 TSPO-PET findings with spatial learning performance and Aβ accumulation. (**A**) Water maze (WM) performance index (principal component analysis, PCA, higher score means better performance) in the comparison of PS2APP mice after five months pioglitazone treatment and their vehicle controls. (**B**) Correlation between the longitudinal change of TSPO-PET in PS2APP mice and the water maze performance index. (**C**) Water maze performance index in the comparison of *App^NL-G-F^* mice after five months pioglitazone treatment and their vehicle controls. (**D**) Correlation between the longitudinal change of TSPO-PET in *App^NL-G-F^* mice and the water maze performance index. (**E**) Aβ-PET rate of change (Δ SUVR) in the comparison of PS2APP mice after five months pioglitazone treatment and their vehicle controls ^16^. (**F**) Correlation between the TSPO-PET rate of change in PS2APP mice and the Aβ-PET rate of change. (**G**) Aβ-PET rate of change (Δ SUVR) in the comparison of *App^NL-G-F^* mice after five months pioglitazone treatment and their vehicle controls ^16^. (**H**) Correlation between the TSPO-PET rate of change in *App^NL-G-F^* mice and the Aβ-PET rate of change. P-values of the group comparisons derive from an unpaired two-tailed t-test. R- and P-values of the correlation analyses derive from a Pearson correlation. PS2APP pioglitazone n=13, PS2APP vehicle n=10, *App^NL-G-F^* pioglitazone n=15, *App^NL-G-F^* vehicle n=15. AUC = area under the curve.

### ^18^F-GE180 TSPO-PET signal reflects activated microglia

Finally, we set about to elucidate the molecular source of the TSPO-PET signal. Earlier studies have already validated *in vivo* TSPO-PET as a microglial marker relative to immunohistochemistry *ex vivo* ^10, 11^ and we have elsewhere demonstrated that PPARγ-related modulation of microglia can be detected by terminal immunohistochemistry in these mouse models ^16^. However, the molecular and cellular correlates of altered TSPO expression during pharmacological PPARγ stimulation remained unclear. To establish this relationship, we performed an immunohistochemical validation of TSPO-PET in subpopulations of all study groups using antibodies against a general marker of microglia (Iba-1) and a specific marker of microglial activation (CD68).

Iba-1 (R = 0.790, p < 0.0001, **Figure 5A**) and CD68 (R = 0.952, p < 0.0001, **Figure 5B**) immunohistochemistry results correlated highly with TSPO-PET binding *in vivo*. Importantly, we saw a stronger association between TSPO-PET with CD68 labelling, which we attribute to the lesser differentiation of Iba-1 immunohistochemistry for treated *App^NL-G-F^* and PS2APP. Indeed, Iba-1 immunohistochemistry did not differentiate between treated *App^NL-G-F^* and treated PS2APP mice. The lacking differentiation of pioglitazone-treated PS2APP and *App^NL-G-F^* mice by Iba-1 immunohistochemistry was also discernible at the individual mouse level (**Figure 5C**). This indicated that PPARγ stimulation specifically reduced the activation status of microglia and that TSPO-PET predominantly tracked activated microglia.

**Figure 5.**
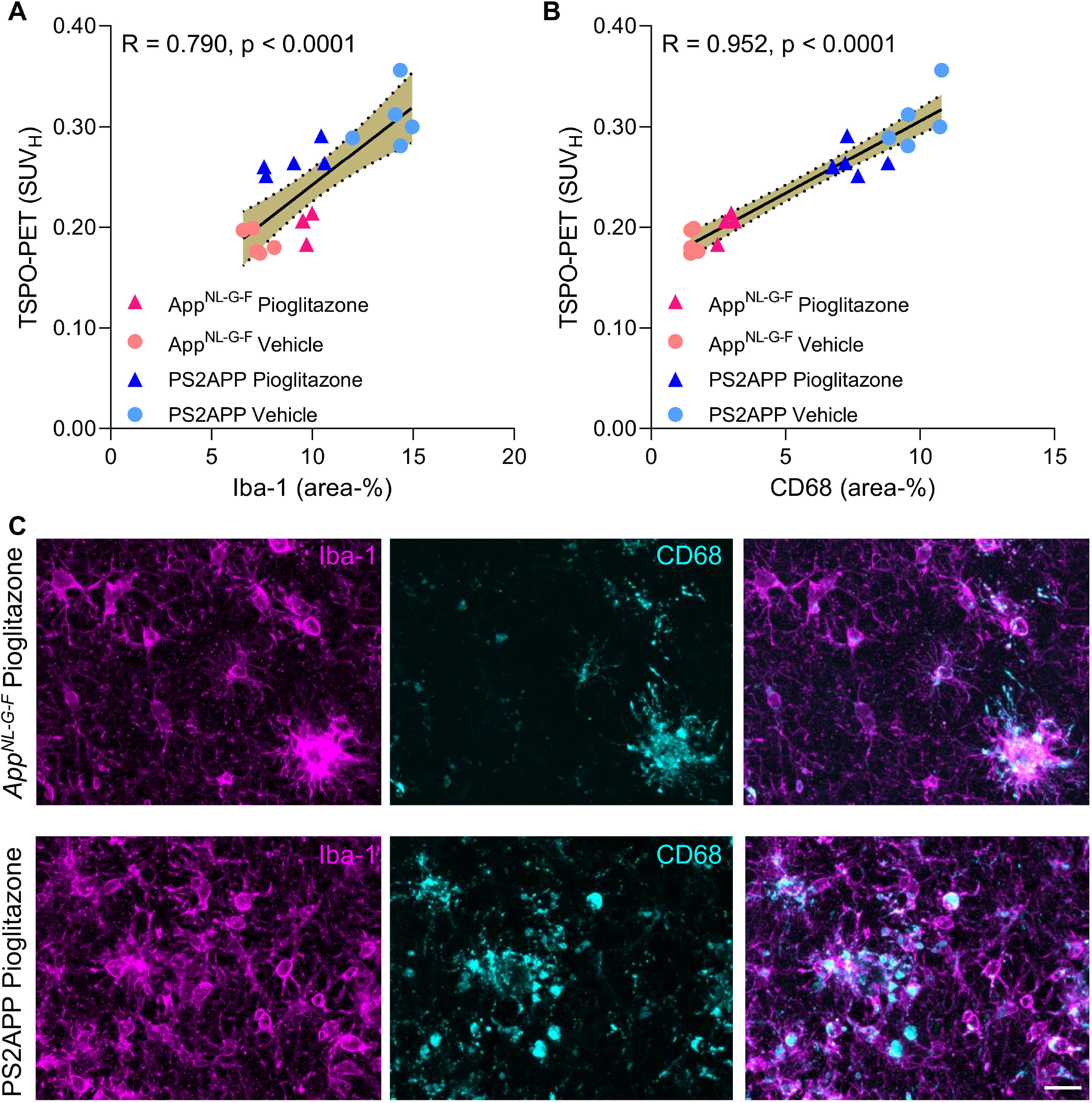
(**A, B**) Correlation analysis between immunohistochemistry markers of microglial activation and ^18^F-GE180 TSPO-PET at the terminal time-point. *App^NL-G-F^* pioglitazone n=4, *App^NL-G-F^* n=5, PS2APP pioglitazone n=5, PS2APP vehicle n=5. Error bands represent the 95% confidence intervals. SUV_H_ = standardized uptake value including myocardium correction. R = Pearson’s coefficient of correlation. (**C**) Representative immunohistochemistry images of treated *App^NL-G-F^* and PS2APP mice, indicating a similar Iba-1 area coverage but a higher CD68 area coverage of PS2APP mice compared to *App^NL-G-F^*. The differentiation by CD68 fitted to corresponding TSPO-PET images, clearly showing an elevated TSPO-PET signal in PS2APP mice compared to *App^NL-G-F^*. Scale bar = 20 μm

## Discussion

In this longitudinal study, we investigated serial TSPO-PET imaging as a tool for monitoring of chronic immunomodulation in two distinct mouse models of amyloidosis. Here, PET with the TSPO ligand ^18^F-GE180 sensitively detected changes of microglial activity upon PPARγ stimulation. Furthermore, we discovered an important sex difference in treatment response. Pre-therapeutic TSPO-PET measures supported a prediction of treatment responses across mouse models and sexes, thus indicating that individual differences in baseline TSPO expression set the stage for effects of immunomodulation. Immunohistochemistry confirmed that TSPO-PET specifically measured activated microglia in the present models.

Our results prove that PET with the TSPO ligand ^18^F-GE-180 can sensitively monitor pioglitazone-induced changes of microglial activity during chronic treatment of AD-model mice. This finding is important given that an earlier PET study using the less avid TSPO ligand ^11^C-(*R*)-PK11195 failed to detect treatment-induced changes in the TSPO-PET signal during chronic pioglitazone administration in APPPS1 mice ^19^. Nonetheless, the ^11^C-(*R*)-PK11195 methodology was sufficiently sensitive to detect microglial activation in transgenic versus wild-type mice. We note that a head-to-head comparison with an equal treatment setting would be required to draw robust conclusions on the superiority of one tracer over the other. However, earlier studies support our present findings of excellent sensitivity for ^18^F-GE-180 TSPO-PET, since the tracer outperformed ^11^C-(R)-PK11195 in a preclinical head-to-head comparison after lipopolysaccharide challenge ^20^ and revealed higher specific binding *in vivo* when compared to ^11^C-PBR23 in a human blocking study ^21, 22^. Importantly, we successfully measured treatment effects on microglial activation by TSPO-PET in two distinct Aβ mouse models. Here, our use of *App^NL-G-F^* mice ^13^ provided evidence that TSPO expression is altered by PPARγ stimulation in mice without overexpression of APP. In line with our data, ^18^F-GE-180 PET also enabled the detection of reduced microglial activation during neurotrophin receptor modulation by LM11A-31 ^9^. In brief, our PET monitoring of pharmacological PPARγ stimulation clarified that the direct modulation of microglial activity can be captured *in vivo*.

The main finding of our study is that pre-therapeutic and serial TSPO-PET recordings in our chronic pioglitazone treatment paradigm allowed predicting individual treatment responses. First, mice with high microglial activation at baseline showed stronger treatment effects, which was underpinned by a strong association between high baseline TSPO-PET quantitation and slower increase rates of TSPO-PET signal during the five months of PPARγ stimulation. Thus, TSPO imaging of microglial activation can serve as a powerful translational tool ^23^, allowing for individual predictive response stratification before or during immunomodulation in the context of precision medicine. Second, the magnitude of microglial activation during the treatment period had close associations with changes in fibrillar Aβ pathology of both models and with spatial learning performance of PS2APP mice. The present finding of stronger increases of fibrillar Aβ in mice with low baseline microglial activity is entirely in line with our translational study in mice with amyloidosis and AD patients ^18^. Thus, the present results strengthen the hypothesis that activated microglia mediate the clearance of excess fibrillar Aβ. Interpretation of the observed association between low microglial activation and better spatial learning performance calls for some subtlety. Although a sufficient microglial response seems important to maintain brain function in therapy-naive AD model mice ^15, 24^, the suppression of microglial activation by PPARγ stimulation was directly correlated with better spatial learning performance in the current study. Thus, we suppose that PPARγ stimulation shifted the already activated microglia (i.e., in mice with high TSPO-PET levels at baseline) towards a more pronounced neuroprotective function. Proving this conjecture might call for a more rigorous discrimination of the M1/M2 phenotypic characteristics. Nonetheless, present data definitely substantiate that microglia play a major role in the histological and behavioral consequences of cerebral amyloidosis in mice.

Interestingly, we observed a pronounced sex effect on the pioglitazone treatment response in *App^NL-G-F^* mice. Vehicle-treated female *App^NL-G-F^* mice showed the previously reported stronger increase of TSPO expression when compared to their male littermates ^12^, but PPARγ stimulation rectified this increase in females while exacerbating the course of neuroinflammation in males. This finding is of remarkable translation significance, since some pioglitazone studies used only female mice ^25^ or did not declare the sex of mice ^26^. Thus, potential sex effects of PPARγ stimulation might have been missed in these studies. Furthermore, this sex effect merits attention consideration in planning human studies, since levels of sex hormones could impact upon microglial modulation ^27^. On the other hand, our parallel detailed analysis of amyloid aggregation during chronic PPARγ stimulation in this cohort did not show relevant sex differences in the rate of increase in Aβ PET signal ^16^. Still, this fits with our previously reported dependency of the Aβ-PET rate of change on microglial activity in AD model mice ^18^, since the present pioglitazone treatment in male and female *App^NL-G-F^* mice resulted in similar microglia activation levels at the end of the study.

We initiated PPARγ therapy at ages manifesting an early phase of limited fibrillar amyloidosis in both mouse models, thus emulating an early but detectable stage of the human AD continuum ^28^. In consideration of emerging plasma biomarkers for AD pathology ^29^, novel treatments for AD shall likely be initiated at a comparable disease stage in future clinical studies. Thus, we focused our intervention monitoring on the phase of amyloid aggregation, which revealed the greatest therapeutic response in AD model mice with a seemingly more aggressive microglial activation during early amyloid build-up. Thus, insofar as PET, cerebrospinal fluid, or plasma biomarkers of microglial activation could serve for treatment stratification in early AD with positive Aβ-status, we foresee opportunities emerging for personalized precision medicine. The major drawback of current mouse AD models is the missing conversion of a sole Aβ-positive stage (A+T-) to combined Aβ/tau-positivity (A+T+). Although the recent literature describes novel combinations of Aβ/tau gene modification ^30, 31^, these models still do not present a breakthrough in better mimicking human AD. Conceivably, cortical tau seeding in an Aβ mouse model might yield a more AD-like model of tauopathy, but such models are not yet ready for large scaled testing of drugs ^32^. Thus, we note as a limitation of the present study that we were unable to investigate effects of pioglitazone on conversion to tau-positivity or during subsequent tau spreading. As a consequence, we cannot predict the efficacy of chronic PPARγ stimulation on the tauopathy encountered at late stages of AD.

The molecular sources of the TSPO-PET signal in neurodegenerative diseases remained to be fully elucidated ^22^. We undertook a correlation analysis between TSPO-PET and immunohistochemistry endpoints in heterogeneous samples of two mouse models, factoring for age, sex, and presence or absence of immunomodulation. Here, we found that the activated microglial marker CD68 proved to have a much better correlation with TSPO-PET signal. In contrast, Iba-1 immunohistochemistry did not distinguish between PS2APP and *App^NL-G-F^* mice after pioglitazone treatment, although the two groups were clearly separated by TSPO-PET and CD68 immunohistochemistry. Thus, the TSPO-PET signal has a close association with disease-associated microglial activation, which also fits to the strong correlations between CD68 and TSPO-PET reported in Trem2-deficient APPPS1 mice ^11^.

PPARγ receptor agonists represent a rather unspecific drug since PPARγ is involved in various pathways in addition of peroxisome activation, notably including glucose metabolism and insulin sensitization ^33^. We selected pioglitazone for immunomodulation of microglial activity in AD mouse models as the effects of this drug are well understood. Nonetheless, more specific drugs like NLRP3 regulators ^34^ could enable a more direct targeting of the inflammasome in neurodegenerative diseases. Optimization of immunomodulator strategies could potentially improve their effectiveness and reduce their side effects, whereupon our present TSPO-PET imaging paradigm could be readily transferable to other drugs, so long as they target activated microglia. Ultimately, specific radioligands for different microglia phenotypes ^35^ could enhance monitoring of immunomodulation *in vivo*.

## Conclusion

TSPO-PET serves as a sensitive biomarker for *in vivo* monitoring of immunomodulation in mouse AD models. Pre-therapeutic assessment of microglial activation in individual mice predicted their response to immunomodulation therapy, indicating that a biomarker of microglial activation could serve for responder stratification. There were pronounced sex differences in the responses to PPARγ stimulation effects *in vivo*. The observed heterogeneity of treatment responses in mice with equal genetic background calls for special consideration in the design of biomarker studies assessing effects of immunomodulation on microglial activation in translational trials in AD patients.

## Material and Methods

### Study design

All experiments were performed in compliance with the National Guidelines for Animal Protection, Germany, with approval of the local animal care committee of the Government of Oberbayern (Regierung Oberbayern) and overseen by a veterinarian. The experiments also complied with the ARRIVE guidelines and were carried out in accordance with the U.K. Animals (Scientific Procedures) Act, 1986 and associated guidelines, EU Directive 2010/63/EU for animal experiments. The chronic treatment study was performed in two different Aβ mouse models and a longitudinal PET imaging design was applied in both cohorts. Female PS2APP and wild-type mice had their baseline assessment at eight months of age and had follow-up PET imaging at 9.5, 11.5 and 13 months of age. Female and male *App^NL-G-F^* mice had their baseline assessment at five months of age and received follow-up PET imaging at 7.5 and 10 months of age. Cage randomization to pioglitazone treatment or control chow (vehicle) groups was initiated after the baseline PET scans, and treatments continued until after the terminal behavioural assessments. After recovering from the final PET scan, mice were transferred to the behavioral facility and rested for one week before initiation of Morris water maze (WM) testing of spatial learning. One week after the behavioural tests, mice were deeply anaesthetized prior to transcardial perfusion and fixation with 4% paraformaldehyde. We then harvested and processed the brains for immunohistochemical and biochemical analyses (randomized hemispheres). Group comparisons of longitudinal Aβ-PET monitoring and detailed Aβ analyses by immunohistochemistry and biochemistry of the same cohort are reported in a separate manuscript ^16^. Shared datapoints between both manuscripts are indicated and cited.

### Animal Models and Statistical Power Analysis

The transgenic B6.PS2APP (line B6.152H) is homozygous for human presenilin (PS) 2, the N141I mutation, and the human amyloid precursor protein (APP) K670N/M671L mutations ^36^. Homozygous B6.PS2APP mice show first appearance of plaques in the cerebral cortex and hippocampus at 5–6 months of age ^37^. The knock-in mouse model *App^NL-G-F^* carries a mutant APP gene encoding the humanized Aβ sequence (G601R, F606Y, and R609H) with three pathogenic mutations, namely Swedish (KM595/596NL), Beyreuther/Iberian (I641F), and Arctic (E618G). Homozygotic *App^NL-G-F^* mice progressively exhibit widespread Aβ accumulation from two months of age ^13, 38^. Both transgenic models were generated on a C57Bl/6 background which also served for wild-type controls.

Required sample sizes were calculated by G*power (V3.1.9.2, Kiel, Germany), based on assumptions for a type I error α=0.05 and a power of 0.8 for group comparisons. A drop-out rate of 10% per time-point was assumed and a treatment effect causing 5% change in the PET signal was considered significant. Estimations were based on PET measures in previous investigations with the same mouse models ^10, 39^. Calculated sample sizes at baseline were n=14 for PS2APP, n=8 for wild-type, and n=9 per sex for *App^NL-G-F^*.

### PET Imaging

For all PET procedures, radiochemistry, data acquisition, and image pre-processing were conducted according to an established, standardized protocol ^40^. In brief, ^18^F-GE-180 TSPO-PET recordings (average dose: 11.5 ± 2.2 MBq) with an emission window of 60–90 min after injection were performed for assessment of cerebral TSPO expression. Aβ-PET recordings (^18^F-florbetaben average dose: 12.2 ± 2.0 MBq) with an emission window of 30–60 min after injection were obtained to measure fibrillar cerebral amyloidosis, as reported elsewhere ^16^. Isoflurane anesthesia was induced before tracer injection and maintained to the end of the imaging time window. All image analyses were performed using PMOD (version 3.5; PMOD technologies, Basel, Switzerland). Static 30-60 min (Aβ-PET) and 60-90 min (TSPO-PET) datasets were co-registered to tracer specific templates (genotype specific) by a manual rigid-body transformation (TX_rigid_) ^40^. In the second step, a reader-independent affine coregistration to the tracer-specific template was performed ^40^. Here, the initial manually fused images were further normalized by non-linear brain normalization (TX_BN_) via the PMOD brain normalization tool (equal modality; smoothing by 0.6 mm; nonlinear warping; 16 iterations; frequency cutoff 3; regularization 1.0; no thresholding). The concatenation of TX_rigid_ and TX_BN_ was then used to obtain optimal resampling with a minimum of interpolation. Normalization of injected radioactivity was performed by the previously validated myocardium correction method ^41^ for TSPO-PET and by previously established white matter^40^ (PS2APP) and periaqueductal grey ^39^ (*App^NL-G-F^*) reference regions for Aβ-PET. TSPO- and Aβ-PET estimates (per time-point and rate of change) deriving from the same neocortical target VOI (15 mm^3^) were extracted and compared between treatment and vehicle groups as well as between transgenic mice and wild-type controls by mixed linear models. The TSPO-PET z-score of each individual transgenic mouse at each time-point was calculated by subtraction of the mean TSPO-PET value of vehicle treated age-matched wild-type mice and division by the standard deviation of wild-type mice (z-score = [mean_TG_ – mean_WT-Veh_]/SD_WT-Veh_). The z-score deviation per time was defined as a TSPO-PET AUC ^15^ and served as an index for microglial activation during the observation time period. For the association analysis between baseline TSPO-PET and changes of TSPO-PET over time (Δ z-score = rate of change), we additionally extracted VOIs from the Mirrione atlas ^42^ to allow evaluation of multiple brain regions. The large cortex VOI of the atlas was split into motor/sensory, auditory/visual and entorhinal/piriform cortices to allow evaluation within compartments of functional similarity. A false discovery rate correction for multiple comparisons was applied.

### Water maze

Two slightly different Morris water maze tasks were applied due to facility changes between the investigation of PS2APP and *App^NL-G-F^* cohorts. We used a principal component analysis of the standard read outs of each water maze task to generate a robust read-out for correlation analyses ^43^. Thus, one quantitative index of water maze performance per mouse was calculated via dimension reduction and correlated with PET imaging. The experimenter was blind according to the phenotype of the animals. Water maze results were also used as an endpoint in the dedicated manuscript on Aβ-PET in both mouse models ^16^.

PS2APP and age-matched wild-type mice were subjected to a modified water maze task as described previously ^15, 44–46^ yielding escape latency, distance to the correct platform and correct choice of the platform as read outs. Mice had to distinguish between two visible platforms, one of which was weighted in such a manner that it would float when the mouse climbed on (correct choice), while the other would sink (wrong choice). The correct platform was always located at the same spot in the maze, while the wrong platform as well as the site from which the mice were released into the maze were varied in a pseudorandom fashion. Visual cues on the walls of the laboratory provided orientation. Trials were terminated if the mouse had failed to reach one of the platforms within 30 sec (error of omission). In this case, or in case of a wrong choice, the experimenter placed the mouse on the correct platform. After a three-day handling period, water maze training was performed on five consecutive days, with five trials per day, which were conducted 2-4 minutes apart. Memory performance was assessed by measuring the escape latency at each day of training and by the travelled distance at the last training day. For measuring escape latency, we calculated the summed average time of all trials from the start point to attaining one of the platforms. On the sixth day, the correct platform was placed in the opposite quadrant of the maze to confirm that the mice indeed used spatial cues rather than rule-based learning to find it. Trials were filmed with a video camera and the swimming trace was extracted using custom written LabView software (National Instruments).

*App^NL-G-F^* mice and 14 age- and sex-matched wild-type mice underwent a common Morris water maze test, which was performed according to a standard protocol with small adjustments ^47^ as previously described ^39^. In brief, the first day was used for acclimatization with a visible platform (five minutes per mouse). The mice then underwent five training days where each mouse had to perform four trials per day with the platform visible at the first training day and the platform hidden under water for all other training days. The test day was set by only one trial with complete removal of the platform. The maximum trial length on all training and test days was set to a maximum of 70 seconds. The video tracking software EthoVision^®^ XT (Noldus) was used for analyses of escape latency, the platform frequency and attendance in the platform quadrant at the probe trial.

The principal component of the water maze test scores was extracted from three spatial learning readouts (PS2APP: escape latency, distance, platform choice; *App^NL-G-F^*: escape latency, frequency to platform, time spent in platform quadrant) using SPSS 26 statistics (IBM Deutschland GmbH, Ehningen, Germany). Prior to the PCA, the linear relationship of the data was tested by a correlation matrix and items with a correlation coefficient < 0.3 were discarded. The Kaiser-Meyer-Olkin (KMO) measure and Bartlett’s test of sphericity were used to test for sampling adequacy and suitability for data reduction. Components with an Eigenvalue > 1.0 were extracted and a varimax rotation was selected.

### Immunohistochemistry

Iba-1 and CD68 immunohistochemistry was performed as described previously ^39, 48^ and the group comparisons between treatment and vehicle groups are reported in the accompanying manuscript ^16^. Correlation analyses were performed between TSPO-PET and Iba-1/CD68 quantitation. n=4-5 PS2APP and *App^NL-G-F^* mice per treatment and vehicle groups with a successful TSPO-PET scan prior to immunohistochemistry were subjected to this analysis. In brief, we performed a standard free-floating immunofluorescence protocol with cortex areas matching the PET brain regions of interest. As previously described, perfusion-fixed 50-μm thick brain sections were rinsed either overnight or for 48 h in PBS with 0.2% Triton X-100 containing one of the following primary antibodies: rabbit monoclonal Iba-1 (1:500. Wako: 19-19741), or rat monoclonal CD68 (1:500. Bio-rad: MCA1857). After washing in PBS, sections were then incubated in a combination of three secondary antibodies (Alexa 488 goat antirabbit, Alexa 594 goat anti-mouse). A detailed analysis of Aβ-plaques (methoxy-X04 and NAB223) of this cohort is reported in the accompanying manuscript ^16^.

### Statistics

Group differences (i.e. between treatment groups or sexes) in TSPO-PET trajectories over time were determined using linear mixed models using the lmer package in the R statistical software, including a random intercept. Note that we selected models including either linear or quadratic time effects based on best model fit (i.e. lower Akaike Information Criterion for better model fit).

Association analyses were performed between PET, water maze, and immunohistochemistry scores. Pearson’s coefficient of correlation (R) was calculated after confirming normal distribution by a Kolmogorov-Smirnow test. Correlation analysis was performed between TSPO-PET baseline (z-score) and the rate of change of TSPO-PET (Δ z-score). This analysis was performed in the cortical target region and in a separate analysis of the full Mirrione atlas set of VOIs ^42^. False discovery rate correction was applied for the multi-region analysis. The rate of change of TSPO-PET (Δ z-score) was correlated with the principal component of the water maze task to investigate potential associations of the PPARγ stimulation treatment effect with spatial learning performance. The index of microglial activity during a certain time-period (AUC) was correlated with the Aβ-PET rate of change to test the hypothesis of Aβ removal by activated microglia ^18^. Immunohistochemistry quantification (Iba-1 and CD68) in the cortex was correlated with the cortical TSPO-PET signal of the terminal time-point.

## Acknowledgements

The study was supported by the *FöFoLe* Program of the Faculty of Medicine of the Ludwig Maximilian University, Munich (grant to M.B.). This work was funded by the Deutsche Forschungsgemeinschaft (DFG, German Research Foundation) to A.R. and M.B. – project numbers BR4580/1-1/ RO5194/1-1. The work was supported by the Deutsche Forschungsgemeinschaft (DFG, German Research Foundation) under Germany’s Excellence Strategy within the framework of the Munich Cluster for Systems Neurology (EXC 2145 SyNergy – ID 390857198). M.B. was supported by the Alzheimer Forschung Initiative e.V (grant number 19063p). We thank Karin Bormann-Giglmaier and Rosel Oos for excellent technical assistance. We thank Christian Haass for excellent support and supervision of the project. We thank Takashi Saito and Takaomi C. Saido for providing the *App^NL-G-F^* mice. GE Healthcare made GE-180 cassettes available through an early-access model. The authors acknowledge Inglewood Biomedical Editing for professional editing of the manuscript.

